# Cultivation of 146 Clinical Strains of *Nocardia* species

**DOI:** 10.1101/302901

**Authors:** Xiang Y. Han, Arginea J. Holmes, Mohammad A. Golshan

**Affiliations:** Clinical Microbiology Laboratory, the University of Texas M. D. Anderson Cancer Center, Houston, Texas 77030

**Keywords:** *Nocardia*, culture media, decontamination, antimicrobial susceptibility

## Abstract

*Nocardia* species generally grow well on culture media in contrast to limited metabolic activities. Yet there are rare studies that have examined systematically the effect of culture media on the clinical recovery of these Gram-positive branching bacilli, particularly in relation to many recently described or revised *Nocardia* species. We analyzed 146 clinical *Nocardia* strains, of 24 species, recovered at the University of Texas M. D. Anderson Cancer Center from 2002-2018 for their growth on culture media. Among media setups to culture routine bacteria, fungi, and acid-fast bacilli (AFB), the AFB media alone recovered 35 strains as did fungal media with 26 strains and routine media with 10 strains. Seventy five strains were recovered on two and three media setups. On AFB media most *Nocardia nova* strains (24 of 29) grew, significantly more than all other strains did. Likewise, all four strains of *Nocardia mikamii* were cultivated solely on AFB media. Fungal media yielded all six strains of *Nocardia wallacei* that were resistant to tobramycin and/or amikacin; this growth could be explained by the presence of gentamycin in one fungal medium. Specimen source also affected the recovery: more *Nocardia* strains grew on every media setup from normally sterile specimens than from nonsterile specimens. These results suggest that diverse culture media support the growth of many *Nocardia* species. The better recoveries of these organisms from AFB media and fungal media are likely attributable to decontamination of specimens on AFB culture, incorporation of selective antimicrobial agents into the media, and prolonged incubation.

## INTRODUCTION

The Genus *Nocardia* represents a large group of aerobic actinomycetes and currently contains 116 species (www.bacterio.net/Nocardia.html, accessed March 28, 2018). These organisms are Gram-positive bacilli with filamentous and branching morphology, partial acid-fast stain, formation of aerial mycelia in the natural environment and in culture, and resistance to lysozyme (1). There are over 50 medically significant *Nocardia* species, which can cause pulmonary, cutaneous, cerebral, and disseminated infections, mostly in immune-compromised hosts (1–4).

The diagnosis of *Nocardia* infection relies on cultivation of the organism in view of the infrequent occurrence and broad differential diagnosis. It is generally thought that *Nocardia* species grow on most culture media in spite of being fastidious. By contrast, these organisms are relatively inactive in metabolism, which has made it difficult or imprecise to render species identification of clinical isolates based solely on traditional phenotypic methods (5, 6, 7). The increasing uses of molecular methods in the past two decades, such as nucleic acid hybridization techniques, restriction fragment length polymorphism, and gene sequencing analysis have remarkably enhanced the discrimination of various *Nocardia* species in both research and clinical settings (4, 8–15). These technical advances have led to new description and/or taxonomic revision of many *Nocardia* species (2, 3).

Culture studies on the clinical recovery of *Nocardia* species are rare in contrast to the recent progress on the recognition of numerous new species, common practice of molecular methods for species identification, and standardization of susceptibility tests. In this study, we assess the impact of culture media on the recovery of 146 clinical strains of *Nocardia* species for correlation with species and antimicrobial susceptibilities. This is an extension of our recent study on the clinical and microbiologic features of 138 *Nocardia* strains.

## RESULTS

A sum of 146 *Nocardia* strains were analyzed in the study. These strains were isolated from specimens with triple media setups that aimed to recover routine bacteria (C RT), fungi (C Fung), and acid-fast bacilli (C AFB). They included 113 strains from the 2002-2012 dataset (4) and 33 strains from 2013-2018. Not included in this study were 33 other strains that were short of one or two types of culture setup. Some of these strains were recovered from blood cultures, most notably *N. nova* (4), that usually lack initial culture setups for fungi and AFB.

As shown in Table 1, the strains included 24 species to indicate a broad spectrum of clinical recovery of these environmental organisms. No single species was predominant; however, common species included *N. nova, N. cyriacigeorgica, N. farcinica, N. abscessus*, and *N. beijingensis*, together accounting for 96 strains or 65.8% of all. Seven species contained a single strain each.

**Table 1.** Growth of *Nocardia* species on culture media for routine bacteria (C RT), fungi (C Fung), and acid-fast bacilli (C AFB)

| *Nocardia* species | No. strains | No. strains with growth on media |  |  |  |  |  |  |
| --- | --- | --- | --- | --- | --- | --- | --- | --- |
|  |  | All three setups | C RT only | C Fung only | C AFB only | C RT + C Fung | C RT + C AFB | C Fung + C AFB |
| *N. nova* | 29 | 8 |  | 2 | 10 | 3 | 2 | 4 |
| *N. cyriacigeorgica* | 27 | 3 | 4 | 4 | 5 | 5 | 4 | 2 |
| *N. farcinica* | 20 | 7 | 2 | 1 | 5 | 2 | 2 | 1 |
| *N. abscessus* | 11 |  | 1 | 5 | 2 | 1 |  | 2 |
| *N. beijingensis* | 9 | 3 | 1 |  | 1 | 2 |  | 2 |
| *N. veterana* | 7 | 1 |  | 3 | 3 |  |  |  |
| *N. wallacei* | 6 |  |  | 5 |  |  |  | 1 |
| *N. asiatica* | 4 |  |  | 1 | 1 |  | 1 | 1 |
| *N. mikamii* | 4 |  |  |  | 4 |  |  |  |
| *N. pseudobrasiliensis* | 4 | 1 | 1 |  |  | 1 | 1 |  |
| *N. aobensis* | 3 |  |  | 2 |  | 1 |  |  |
| *N. brasiliensis* | 3 |  |  |  |  | 2 | 1 |  |
| *N. carnea* | 3 |  |  |  | 2 |  | 1 |  |
| *N. kruczakiae* | 3 | 1 |  |  |  | 2 |  |  |
| *N. asteroides* | 2 | 1 |  |  |  |  |  | 1 |
| *N. paucivorans* | 2 |  |  |  | 1 |  | 1 |  |
| *N. pneumoniae* | 2 |  |  | 1 |  | 1 |  |  |
| *N. africana* | 1 |  |  | 1 |  |  |  |  |
| *N. araoensis* | 1 |  |  |  |  | 1 |  |  |
| *N. higoensis* | 1 |  |  |  | 1 |  |  |  |
| *N. mexicana* | 1 |  | 1 |  |  |  |  |  |
| *N. otitiscaviarum* | 1 |  |  |  |  |  |  | 1 |
| *N. vinacea* | 1 |  |  | 1 |  |  |  |  |
| *N. yamanashiensis* | 1 | 1 |  |  |  |  |  |  |
| Total | 146 | 26 | 10 | 26 | 35 | 21 | 13 | 15 |

Among the three types of culture setups, C AFB recovered the most *Nocardia* strains (Table 1). There were 35 strains cultivated from C AFB alone, in comparison to 26 strains from C Fung alone and 10 strains from C RT alone. Seventy five strains also grew on two and three culture setups. Together, 89 strains (61.0% of all) grew on C AFB, 88 strains (60.3%) on C Fung, and 70 strains (47.9%) on C RT. Thus, triple setups have helped to cultivate more *Nocardia* strains.

The media setups affected the recovery of three species most. *N. nova* was cultivated mainly on C AFB, with growth of 24 of the 29 strains, significantly more than all other *Nocardia* species combined on these media (65 of 117 strains) (χ^2^ = 7.23, p = 0.0072). Similarly, all four strains of *N. mikamii* were recovered from C AFB alone, whereas only 31 of all rest 142 strains were isolated this way (Fisher’s p = 0.0029). On fungal media grew all six strains of *N. wallacei* with five of them on these media solely. This was different from all other 140 strains in that only 21 were recovered on C Fung alone (p = 0.0007). This culture difference can be explained by the incorporation of gentamycin in the brain heart infusion slant for fungal culture. *N. wallacei*, previously known as *Nocardia asteroides* drug pattern IV (16), is a recently proposed species with a rare feature of resistance to amikacin. As shown in Table 2, such is the case in these *N. wallacei* strains: four of the six were resistant to amikacin, in comparison to only four of the rest 125 tested strains (p = 0.0001). Similarly, all were resistant to tobramycin whereas only 47 of the rest 114 tested strains were this way (p = 0.0063). Gentamycin, amikacin, and tobramycin belong to the class of aminoglycoside with similar antimicrobial activities.

**Table 2.** Susceptibility of *Nocardia wallacei* strains to amikacin and tobramycin

| *Nocardia* species | No. strains with amikacin |  |  | No. strains with tobramycin |  |  |
| --- | --- | --- | --- | --- | --- | --- |
|  | Tested | Resistant | Susceptible | Tested | Resistant | Susceptible |
| *N. wallacei* | 6 | 4 | 2 | 6 | 6 | 0 |
| All others | 125 | 4 | 121 | 114 | 47 | 67 |
| Total | 131 | 8 | 123 | 120 | 53 | 67 |
| Fisher's test | p=0.0001 |  |  | p=0.0063 |  |  |

The impact of specimen source on culture recovery was also examined. During the 16 study years, an estimated 40,000 specimens, from roughly equal numbers of normally sterile and nonsterile sources, were set up for cultures in triples. As shown in Table 3, 49 strains and 97 strains were recovered from normally sterile and nonsterile specimens respectively. Of the strains from sterile sources, 35 strains (71.4%) grew on C RT, 31 strains (63.3%) on C Fung, 39 strains (79.6%) on C AFB, and 18 strains (36.7%) on all media setups. In contrast, of the strains from nonsterile sources, relatively fewer grew on C RT (35 strains, 36.1%) (p = 0.0001), on C AFB (50 strains, 51.5%) (p = 0.0001), and on all media setups (8 strains, 8.2%) (p = 0.0001) whereas C Fung remained similar (57 strains, 58.5%) (p = 0.60, not significant). Thus, sterile specimens favor the recovery of *Nocardia* species, much the same for other microorganisms. The 97 strains of nonsterile sources were all from respiratory tract and recovered mainly from fungal and AFB media. This could be explained by better control of other bacteria, such as incorporation of antimicrobial agents in both setups and decontamination for C AFB, and longer incubation to recover low inoculum and/or slowly growing species.

**Table 3.** Cultivation of *Nocardia* species from normally sterile and nonsterile specimens NS: not significant.

| Specimen source | No. all strains (%) | No. strains (%) with growth on media |  |  |  |
| --- | --- | --- | --- | --- | --- |
|  |  | C RT | C Fung | C AFB | All media |
| Total | 146 (100) | 70 (47.9) | 88 (60.3) | 89 (61.0) | 26 (17.8) |
| Normally sterile | 49 (100) | 35 (71.4) | 31 (63.3) | 39 (79.6) | 18 (36.7) |
| Normally nonsterile | 97 (100) | 35 (36.1) | 57 (58.8) | 50 (51.5) | 8 (8.2) |
| Statistical test | | $\chi^2 = 16.3$, p = 0.0001 | $\chi^2 = 0.276$, p = 0.60, NS | $\chi^2 = 10.8$, p = 0.001 | $\chi^2 = 18.1$, p = 0.0001 |

NS: not significant.

The better recoveries of *N. nova* and *N. mikamii* on C AFB were examined further for explanations. Of the 29 *N. nova* strains, C AFB showed growth of 7 strains from 8 sterile specimens and growth of 17 strains from 21 nonsterile specimens. Thus, from sterile specimens, C AFB recovered similar proportions of *N. nova* (7 of 8, 87.5%) and of other species combined (32 of 41, 78.0%) (p = 1.0). In contrast, from nonsterile (airway) specimens, C AFB recovered most *N. nova* strains as well (17 of 21, 81.0%) but less than half of other species combined (33 of 76, 43.4%) (p = 0.0028). This difference suggested that *N. nova* tolerated the decontamination process. Alternatively or in addition, as the most commonly isolated species (Table 1), *N. nova* might have relatively high organismal density in the way. Thus, *N. nova* is a common and hardy species in the genus. The four *N. mikamii* strains were cultivated from a lung tissue aspirate (sterile), a bronchoalveolar lavage, and two bronchial washings. The lung aspirate strain grew upon 5 days of incubation on C AFB whereas concurrent C Fung and C RT did not yield any microorganism. The remaining strains grew upon decontamination and incubations of 7 to 12 days while concurrent C RT grew normal flora and/or *Staphylococcus aureus* and C Fung had no growth. Thus, decontamination and preference for AFB media likely made the difference.

In view of the supplement of trimethoprim in the MP bottle for C AFB, the susceptibilities of the *N. nova* and *N. mikamii* strains were also examined. All the strains were susceptible to trimethoprim/sulfamethoxazole with minimum inhibitory concentrations ranging from < 0.25/4.8 mg/L to ≤ 1/19 mg/L, well below the resistance breakpoints of 4/76 mg/L set for the drug combination against *Nocardia* (17). Therefore, trimethoprim alone in the liquid medium at 0.5 mg/L hardly affected the growth of *N. nova, N. mikamii*, and likely other *Nocardia* species that also grew on C AFB (Table 1).

Together, the results above suggest that diverse culture media support the growth of many *Nocardia* species. Furthermore, control of other rapidly growing bacteria via decontamination and/or incorporation of selective antimicrobials in the fungal and AFB media combined with longer incubation time improves recovery of these organisms from nonsterile specimens.

## DISCUSSION

Among the 116 *Nocardia* species recognized so far, the environmental niches and geographic distributions are likely diverse despite lack of comprehensive survey. The clinical recovery of 24 *Nocardia* species in this study, nearly half of all medically significant *Nocardia* species, is a reflection of this diversity to some extent. This may be attributable to the warm and wet climate in the greater Houston area, the susceptible and diverse cancer patient population, intense diagnostic procedures and culture effort, relatively large number of strains, and accurate identification of many recently described species.

The diversity of *Nocardia* species is also reflected on their genome blueprints that have been revealed in recent years. For instances, *N. farcinica* has a genome of 6.0 mega-base pairs (Mb) (18) whereas *N. brasiliensis* has 9.4 Mb (19). Even within a species, diversity among strains may be substantial, as shown in a new genome study in which some of our Houston area *N. nova* strains were found to harbor genomes of variable sizes from 7.0 Mb to 8.0 Mb (20). Some pathogenic traits also relate to species: *N. farcinica* has a tendency to infect skin and brain tissues whereas *N. nova* may invade the bloodstream in addition to the more common respiratory infections (4).

It has been our experience that *Nocardia* species are not as fastidious as thought before. Many strains from sterile specimens grew upon 48 hours of incubation on blood or chocolate agar, although detailed data on all strains were difficult to obtain. Surprisingly, the AFB media are the best to recover *Nocardia* species. This result can be explained by the prolonged incubation, decontamination of nonsterile specimens, and/or supplement of antimicrobials in the liquid medium.

Decontamination with NaOH and N-acetyl-L-cysteine controls most rapidly growing bacteria that would otherwise overwhelm the growth of AFB. Lower respiratory tract specimens contain contaminants, yet they are also the most common sources of AFB and *Nocardia*. AFB are resistant to decontamination. *Nocardia* species are partially acid fast due to the presence of mycolic acids in the cell wall. The lengths of these mycolic acids are 44-64 carbons (1, 16, 21, 22), shorter than the length of 60-90 carbons of AFB (1). As demonstrated experimentally earlier (23), decontamination reduces culture recovery of *Nocardia* significantly. Yet, our recovery of 50 strains of various *Nocardia* species, *N. nova* in particular, on AFB media upon decontamination (Table 3) suggests that these organisms, when in clinical specimens, are by and large resistant to the harsh treatment. Thus, decontamination also clears the way for cultivation of *Nocardia* species, in addition to the more common AFB, from nonsterile specimens.

From nonsterile specimens, fungal media recover *Nocardia* species well, better than routine and AFB media (Table 3). This is consistent with incorporation of antimicrobials in the media and longer incubation. Finally, our cultivations of *N. wallacei* and *N. mikamii*, two recent and uncommon species (16, 21), on fungal media and AFB media respectively, have revealed an interesting aspect of these organisms. More future studies on these species may be needed for corroboration and further delineation.

## MATERIALS AND METHODS

### Study setting and patients

The *Nocardia* strains were consecutive (sporadic) isolates from June 2002 to March 2018 at the University of Texas M. D. Anderson Cancer Center, Houston, Texas. The institution has been a comprehensive cancer center with 500 beds in 2002 that increased to 650 beds in 2018. All patients with *Nocardia* isolates had a primary diagnosis of cancer. Approximately two thirds of patients were from greater Houston area and other parts of Texas. Patients from other States and other countries usually stayed in Houston for extended period of time for their cancer care. In view of the environmental origin of *Nocardia*, the climate in Houston included abundant rainfall (around 140 cm per year on average) and warm temperature (monthly means of 12.4°C in the coldest January to 29.1°C in the warmest July and August).

### Cultures setups

Routine cultures (C RT) were set up to recover bacteria from lower respiratory tract specimens (sputa, tracheal aspirates, bronchial washings, broncho-alveolar lavages) and other specimens (wounds, body fluids, tissue aspirates, etc). These cultures included one plate each of sheep blood agar, chocolate agar, MacConkey agar, and colistin nalidixic acid agar for incubation at 35C for at least 48 hours. If a specimen is sterile, setups also included one thioglycolate broth and one trypticase soy broth for incubation of seven days.

Cultures to recover fungi (C Fung) included two plates of Emmons Sabouraud dextrose agar (SDA) and three slants, i.e., SDA, brain heart infusion 10% sheep blood agar with gentamycin and chloramphenicol, and Mycosel agar with chloramphenicol and cycloheximide. The fungal media were incubated at 35°C for up to 28 days.

Cultures for acid-fast bacilli (C AFB) were set up in one Lowenstein-Jansen slant and one BacT/Alert MP liquid bottle (BioMerieux, Durham, NC) and incubated for 8 and 6 weeks respectively. Sterile specimens, such as tissues and body fluids, were inoculated into media directly without processing. Nonsterile specimens from wounds and lower respiratory tract were processed with 1.5% NaOH and 0.5% N-acetyl-L-cysteine (AlphaTec Systems, Vancouver, WA) for 10 to 15 minutes for contamination and concentration. The MP bottle was also supplemented with a cocktail of amphotericin B, azlocillin, nalidixic acid, polymyxin B, trimethoprim (final concentration of ~0.5 mg/L), and vancomycin to inhibit molds and non-AFB. The BacT/Alert instrument detected bacterial growth every 10 minutes. In case a non-AFB grew, the bottle was processed again and re-incubated.

A *Nocardia* isolate was counted as a unique strain if the same species was isolated sequentially from repeat sampling or from different sources of sampling within one admission of a patient.

### Species identification

Initial recognition of *Nocardia* included branching Gram-positive bacilli, positive partial acid-fast stain, and chalky white colonies on culture plates if applicable. These bacilli were further identified by sequencing a portion of the 16S rRNA gene as previously described (4, 24). Briefly, target DNA was extracted from isolated colonies or a positive liquid MP bottle using an extraction buffer (Prepman Ultra, Applied Biosystems Inc., Carlsbad, CA). A polymerase chain reaction was used to amplify the target. For most isolates, a ~600-bp region corresponding to nucleotides 459-1,050 of *N. asteroides* ATCC 19247 (GenBank accession DQ659898) was amplified using two universal bacterial primers 5’TGCCAGCAGCCGCGGTAATAC (forward) and 5’CGCTCGTTGCGGGACTTAACC (reverse). For some isolates whose identity remained ambiguous after analysis of this amplicon, such as those of *N. veterana* and *N. kruczakiae*, additional amplification and sequencing of a more proximal ~600-bp region of the gene was performed. The primers were 5’GCGTGCTTAACACATGCAAGTC (forward) and 5’TCCTCCTGATATCTGCGCATTC (reverse), corresponding to nucleotides 13-659 of *N. asteroides* ATCC 19247 (DQ659898). The two overlapping amplicons covered ~1030-bp of the 16S gene to allow confident identification of nearly all strains to species level in this study. Bidirectional consensus sequences were queried to the NCBI database using the basic local alignment search tool (BLAST) function, and the best identity matches to a reference sequence of a type strain, usually ≥ 99.6%, yielded species identification.

### Data retrieval and analysis

Electronic culture records (Cerner work cards) on every *Nocardia* strain were retrieved and examined. Extracted data included specimen type, culture setups, growth or no growth of *Nocardia*, species identification, and susceptibility to antimicrobial agents. The susceptibility was performed at a reference laboratory according to the guidelines of Clinical and Laboratory Standards Institute (17). Those strains with all culture setups, i.e., C RT, C Fung, and C AFB were subjected to analyses. Correlations with specimen types, species identification, and susceptibility to antimicrobial agents were also performed. When appropriate, statistical analyses were performed by using Fisher’s exact method or χ^2^ test method.

## Acknowledgement

This work was supported in part by the National Institutes of Health grant CA16672 for the DNA Analysis Core Facility at M. D. Anderson. The authors thank the staff at our DNA Analysis Core Facility and Molecular Diagnostic Laboratory for DNA sequencing, the staff of microbiology laboratory for cultures, and the clinical staff for taking care of the patients. The authors declare no conflict of interest.

